# Fine-Scale Population Genetic Structure of Dengue Mosquito Vector, *Aedes aegypti* and its Association to Local Dengue Incidence

**DOI:** 10.1101/561621

**Authors:** Thaddeus M. Carvajal, Kohei Ogishi, Sakiko Yaegeshi, Lara Fides T. Hernandez, Katherine M. Viacrusis, Howell T. Ho, Divina M. Amalin, Kozo Watanabe

**Author notes:** Corresponding author (KW).

## Abstract

Dengue fever is an important arthropod-borne disease which is transmitted by the mosquito vector, *Aedes aegypti*. Vector control programs rely heavily on targeting the mosquito vector in order to stop the disease transmission cycle. Hence, the present study conducted a fine-scale population genetics of *Ae. aegypti* in a highly urbanized and dengue endemic region in the Philippines. Furthermore, the study also explored the correlation of population genetic indices to the local dengue incidence of the region. The genetic diversity and population structure of *Ae. aegypti* populations were analyzed by genotyping 11 microsatellite loci from 526 adult mosquitoes sampled in 21 study areas in Metropolitan Manila. Five genetic indices and its dengue incidence were then correlated using Pearson’s correlation. Results showed low genetic differentiation among mosquito populations indicating high gene flow activity in the region. However, the study also revealed a considerable number of inferred genetic clusters (K=5). The constructed UPGMA dendrogram exhibited close proximity of genetically-similar *Ae. aegypti* mosquito populations that extends in long distances suggesting passive dispersal ability of the mosquito vector. Moreover, a positive and significant correlation was observed between dengue incidence and inbreeding coefficient (*F*is) (r = 0.52, p = 0.02). Overall, the study showed that population genetic structuring can occur in a fine-scale area which consisted notable clustering and extending patterns of genetically-similar mosquito populations. This infers the potential migration ability of *Ae. aegypti* in different locations of the region where specific vector control zones could be carried out to disrupt its dispersal ability. Also, this is the first study that attempted to correlate genetic indices to dengue incidence that could serve as a supplementary index in identifying high dengue risk areas in the future.

**AUTHOR SUMMARY:** Dengue disease puts billions of people worldwide at risk. To mitigate this risk, population genetic studies of its vector, *Aedes aegypti*, are being conducted. The information established from these studies can be utilized to reduce mosquito population and thereby, reduce the opportunity for dengue transmission. In this study, we used microsatellite markers to determine genetic structure and diversity followed by correlation analyses between genetic indices and dengue incidence. Results show a low genetic differentiation among mosquito populations in Metro Manila; it also indicates population genetic structuring in a fine-scale area. This suggest a pattern of migration activity of *Ae. aegpyti* which can be used to mitigate dengue transmission. Moreover, the study also explored in correlating genetic indices and local dengue incidence where it demonstrated significant correlation with the inbreeding coefficient (*F*is). Further investigation is needed on how these genetic indices may be utilized in predicting and identifying high dengue risk areas in endemic areas.

## INTRODUCTION

Dengue disease is the most prevalent mosquito-borne viral infection in tropical and subtropical countries [1] with approximately 2.5 billion people worldwide at risk of contracting the disease [2]. Dengue virus is transmitted primarily to humans by the principal mosquito vector, *Aedes aegypti*. This mosquito species is considered to be the most efficient vector of arboviruses because of its highly adaptive nature to the urban environment [3]. Although a dengue vaccine is available [4], the World Health Organization [2] still recommends disease prevention and control towards the mosquito vector.

Molecular genotyping of the mosquito vector using microsatellites has provided useful insights towards the improvement of mosquito vector control strategies [5,6]. For instance, revealing the gene flow pattern among *Ae. aegypti* populations can be interpreted as the mosquito vector’s dispersal pattern [7,8]. Microsatellites is widely used as the standard molecular marker of choice in population genetic studies of *Ae. aegpyti* [9]. Due to the marker’s high polymorphism, co-dominance, and broad genome distribution [10], it has been deemed suitable for differentiating both macro- and micro-geographic scale mosquito populations [11,12,13].

Despite the many population genetic studies of *Ae. aegypti* using microsatellites worldwide, only a handful of studies have investigated the vector’s genetic structure in a fine-scale area [7,11,12,13,14,15,16]. These fine-scale studies are defined as having sampling points within city boundaries, or villages with geographic distances of less than 50 km. For instance, spatial genetic differentiation across *Ae. aegypti* populations was evident in spatial scales within city boundaries [7,11,12,16], among villages [13,14] and along a street [15]. It has been claimed that genetic divergence in small spatial scales is common in Southeast Asia [11] and could also be attributed to the type of breeding sites [15]. Furthermore, multiple inferred clusters of genetic mosquito populations (K= 3–9) were also detected within these fine-scale areas [11,14,16]. It was suggested that a single house or groups of closely situated houses may act as assembling units in forming these genetic clusters [11].

The information and patterns identified by the population genetic approach can be factored as part of the strategy in reducing the mosquito population, thereby, decreasing the opportunity of dengue transmission. Early fine-scale population genetic studies of *Ae. aegypti* [7,8,11,15] had only demonstrated the degree or magnitude of genetic structuring while recent studies [12,16] concentrated on hypothesis testing such as the role urbanization in genetic divergence [14]. Although the results provided relative insights to our understanding of the mosquito vector, it lacks in demonstrating the clustering or distribution of genetically-similar mosquito populations which can reveal notable patterns for it application in vector control. For instance, in Yuunan Province of China, *Ae. aegypti* populations from border areas of the region are genetically-similar among each other and distinct from its two main cities indicating different invasion and colonization conduits [17]. The findings was presented simply in a dendrogram which can used in creating a sound basis in disrupting possible invasion of this mosquito vector from neighboring countries. In previous fine-scale population genetic studies, such graphical presentation was not illustrated but only described or portrayed in tabular records of pairwise genetic differences (e.g. *F*_ST_, Nei’s genetic distance). Illustrating how geographical locations are genetically-similar or distinct may demonstrate the migration activity of *Ae. aegypti* and the extent of its spatial distribution patterns in fine-scale areas.

Surveillance of the immature or adult stages of *Ae. aegypti* has led to the conception of vector indices (e.g. Container, House, Breteau, Pupal or Adult indices) which can be utilized as a potential predictor of local dengue epidemiology [18]. However, frequent or consistent sampling of immature or adult stages of the mosquito vector has proven to be laborious and cost intensive [19]. Innovative and alternative avenues are now being explored in determining the population size of *Ae. aegypti* such as the development of non-powered passive adult traps [20] and utilizing container-inhabiting mosquito simulation approach [21,22]. One avenue that has not been explored is the application of genetic indices that characterizes the mosquito vector population. For example, population genetics can estimate the effective population size *(Ne)* of the mosquito vector which is related to its consensus size [23]. With several population genetic studies that have reported the estimated population sizes of local *Ae. aegypti* populations [9,13,24], no study had associated *Ne* or other genetic indices to the local dengue incidence. It was demonstrated that sampling 25-30 individuals per local area is suitable for microsatellite-based population genetic studies [25] where genetic analyses can determine mosquito population size by conducting specific sampling episodes as compared to frequent or regular mosquito collection surveillance.

Therefore, the present study has two objectives. First, it determines the population genetic structure of *Ae. aegypti* within a fine-scale area and, second, it explores the correlation of population genetic indices to the local dengue incidence. The results can provide a basis of creating new and innovative approaches in controlling this mosquito vector in highly urbanized or endemic areas in the Philippines.

## METHODS

### Study area and Mosquito sampling

Metropolitan Manila, a highly urbanized area, is the National Capital Region (NCR) of the Philippines with an area of 636 km^2^. It is located at the eastern shore of Manila Bay in Southwestern Luzon (14°50’ N Latitude, 121°E Longitude), Philippines, Southeast Asia. It is composed of 16 cities and 1 municipality with a total population of 12,877,253 [26]. This area is the most urbanized region in the Philippines being the center of the national government, economy, education and culture, country’s leading business center, largest manufacturing location and principal port for importation and exportation [27].

In this study, Metropolitan Manila was divided into 21 study areas (Figure 1 and Table S1). Initially, study areas are delineated per city which represents at least one per city. However, some cities comprised of either a large or small land sized area, thus designating more study areas. For example, the largest city, Quezon City, in the region was divided into 5 study areas based on its district boundaries. In order to standardize the land size covered by each study area, we merged two small neighboring cities, San Juan and Mandaluyong. The map layer of the administrative city boundaries of Metropolitan Manila was obtained from the Philippine Geographic Information System (GIS) Data Clearinghouse for further analysis [28].

**Figure 1.**
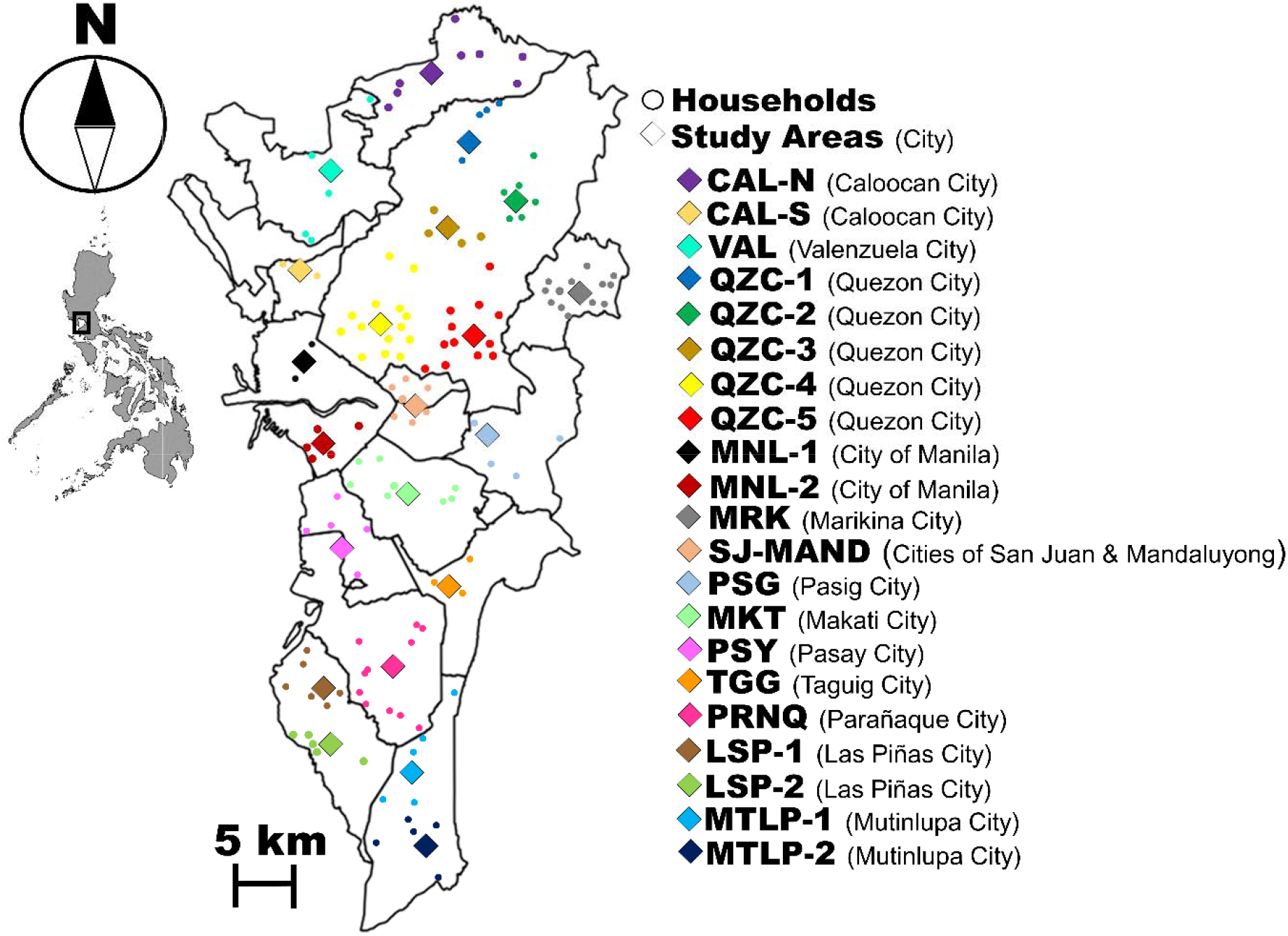
Geographic midpoints of *Ae. aegypti* study areas (◊) with its corresponding household sites (○) in Metropolitan Manila. Details of each study area can be seen in Supplementary Table S1.

Households on each study area were selected based on voluntary informed consent in collecting adult *Ae. aegypti* mosquitoes inside their premises. The number of households per study area ranged from 2 – 14 with an average of 6 households and a total of 134 households. The maximum distance among households within study areas ranged from 1.39 km to 6.17 km. Since the households are widely dispersed within each study area, we calculated the geographical midpoint (http://www.geomidpoint.com/) to assign a single georeferenced location for each study area in subsequent genetic analysis. The distance among study areas (midpoints) ranged from 2.85 – 39.66 km.

Collection of *Ae. aegypti* mosquitoes was done by installing a commercially available mosquito UV Light Trap (Jocanima ©) in each household’s outdoor premises for 3-5 days. Collected adult mosquito individuals were sorted, then identified accordingly based on the pictorial keys of Rueda et al. [29] and preserved in 99% ethanol. Majority of population genetic studies in *Ae. aegypti* have either used only larval or reared larval to adult samples. For this reason, this could lead to a potential bias in estimating population genetic parameters due to the sampling of full sibling mosquito larvae [30]. This was evidence in the population genetic structure of amphibian larval samples that led to inaccurate estimate of differentiation among populations when compared to adult samples [31,32]. As such, this study targets adult *Ae. aegypti* samples than conventional egg, or larval samples to prevent collecting mosquito full siblings. From a collection period of May 2014 until January 2015, a total of 526 adult *Ae. aegypti* were collected and ranging from 12 to 42 individuals per study area (Table S1).

### DNA Extraction and microsatellite genotyping

The total genomic DNA of individual mosquito sample was extracted using the QIAGEN Blood and Tissue DNEasy Kit following the modified protocol of Crane [33]. We identified 11 microsatellites from Slotman et al. [34] and Chambers et al. [10] for genotyping and grouped them accordingly into four sets for multiplex PCR (Table S2). Each set consisted fluorescent labeled forward primers with different annealing temperatures during the amplification process. Generally, each set composed of 1.2 μL of 10X buffer (TAKARA), 0.8 μL of 25 mM MgCl_2_, 1.6 μL of 10 mM of each dNTPs, 0.6 μL of 10 μM forward and reverse primers and 0.08 μL of 5.0U/ μL of Taq DNA polymerase (TAKARA), 1.5 μl of 10% Dimethyl sulfoxide (DMSO) and 1 μl of template DNA consisting a final volume of 10 μl. Thermocycle conditions are as follows: initial denaturation step of 94 □C for 5 minutes, denaturation step of 94 □C for 30 seconds, annealing step with temperature and duration (in seconds) of each primer set as indicated in Table S2, extension step of 72 □C for 30 seconds following 35 cycles and a final incubation step of 72 □C for 5 minutes. PCR amplicons were checked by electrophoresis in 3% agarose gels stained with Midori Green (Nippon Genetics) and visualized under UV light using the Chemidoc XRS Chemiluminescent Gel Documentation Cabinet (BIO-RAD). Prior to fragment size analyses, multiplex PCR products were diluted in 1/15 water and then pooled together. 1ul of each diluted pool were added with 0.5 μl of GS 500 Liz Internal Size StandardTM (Applied Biosystems, USA) and HD formamide for a total volume of 20 μl. Fragment analysis of the amplified products were done using ABI 3500 Genetic Analyzer (Life Technologies) while genotyping is done using GeneMapper (Applied Biosystems). Microsatellite data were checked for error and the presence of null alleles with MICROCHECKER [35].

The exact Hardy-Weinberg equilibrium (HWE) test and estimations of the Linkage disequilibrium (LD) among all pairs of loci were conducted using GENEPOP v4.2.1 [36,37]. Significance levels for multiple testing were corrected using the Bonferroni procedure. The number of alleles, allelic richness and private alleles were calculated using HPRARE [38,39]. Observed heterozygosity (*Ho*), expected heterozygosity (*He*), inbreeding coefficients (Fis) were calculated using the Genetic Analysis in Excel (GenAlEx) version 6.3 [40]. To assess the magnitude of genetic differentiation among sites, pairwise F_ST_ values were calculated using Arlequin v3.5.1.3 [41] with 10,000 permutations. Pairwise gene flow estimates (*Nm*) among sites were manually calculated using the formula of Slatkin and Barton [42] from the calculated pairwise FST. A dendrogram was constructed based on the pairwise *F_ST_* using the unweighted pair group with arithmetic mean (UPGMA) in *fastcluster* package [43] and the optimal number of clusters were determined using the cindex index in the *NbClust* package [44] from the R program [45].

### Genetic Structure

The number of genetic clusters (K) was inferred using the Bayesian approach in the software STRUCTURE v2.3.2 [46]. The admixture model was utilized where its alpha value was allowed to vary, and independent allele frequencies was set at lambda equals to one. Twenty (20) independent runs were performed for each value of K (1 – 15) with a burn-in phase of 200,000 iterations followed by 600,000 replications. Structure Harvester v0.6.93 [47] was used to determine the most likely number of clusters by calculating ΔK [48]. Moreover, the software program CLUMPP v1.1.2 [49] was used to summarize the results from STRUCTURE and visualized using the program DISTRUCT v1.1 [50].

### Isolation by Distance and Spatial Autocorrelation

Pairwise geographic distances (km) among study areas and households were calculated using the Vincenty’s formulae [51] on Microsoft Excel 2016. To test isolation by distance (IBD), pairwise *F_ST_* and geographic distance (km) among study areas were examined using Mantel’s test of correlation with 10,000 permutations. Spatial autocorrelation was performed using pairwise Nei’s genetic distance among mosquito individuals and geographic distance (km) among households with 10,000 permutations and Bootstrap replications. Results of the permutation were considered significant at the 5% level. In this analysis, a correlogram was produced with 45 distance classes at 1km interval. Both analyses yielded a correlation coefficient of the two data matrices ranging from −1 to +1, with a test for a significant relationship by random permutation. All analyses were performed using GenAlEx version 6.3 [40].

### Correlation Analysis between Genetic indices and Dengue incidence

In order to calculate the dengue incidence of each study area, reported dengue cases per village (*baranggay*) in 2014 were obtained from the National Epidemiology Center of the Department of Health, Philippines while the population census per village were acquired from the Philippine Statistics Authority agency (www.psa.gov.ph). Calculation of dengue incidence was performed by dividing the number of cases to the total population size for a given year multiplied by a factor of 1,000. Pearson’s correlation coefficient was calculated based from the computed dengue incidence and the selected population genetic indices namely; Allelic richness, Private Allelic Richness, Observed heterozygosity, Inbreeding coefficient and the effective population size. The correlation analysis was performed using the *stats* package of the R program version 3.3.5 [45].

## RESULTS

### Genetic Diversity

We observed a total of 113 alleles across 11 microsatellite loci in 21 study areas from Metropolitan Manila (Table S3). The number of alleles per loci ranged from 3 (F06) to 19 (B07) with an average of 10.25 alleles per loci, suggesting that the chosen microsatellites markers are highly polymorphic (Table S4). Null alleles were present in 4 loci (M313, AC4, AG7 and H08) and the null allele frequency ranges from 0.00 – 0.33 in all loci (Table S5). For the 231 tests of HWE of each locus per study area, 91 tests showed statistically significant deviation (p < 0.05) where 72 of these significant deviations indicated He > Ho, suggesting heterozygosity deficits (Table S3). The LD test showed a total of 119 of 1155 (10.30%) pairs of loci with significant LD after Bonferroni corrections.

Table 2 shows the summary of the genetic diversity per study area. The mean number of different alleles for all study areas ranged from 3.82 (TGG) to 6.36 (QZC-3) while the mean number of effective alleles for all study areas showed to be 2.74 (QZC-1) to 3.51 (LSP-2). On the other hand, the mean allelic richness ranged from 3.24 (QZC-1) to 3.85 (MRK) for all study areas while the proportion of private alleles ranged from 0.02 (LSP-2) to 0.23 (PSY). Overall, all study areas except for one (MKT) did not conform to Hardy-Weinberg equilibrium expectations (*He > Ho*), indicating heterozygosity deficits and the possibility of inbreeding within each study area. The effective median population size (*Ne*) across all study areas was calculated to be from 6.2 to 4,607 with two study areas estimated as an infinite number.

**Table 2.** Dengue Incidence and Genetic Diversity among 21 *Ae. aegypti* populations based on 11 Microsatellites in Metropolitan Manila, Philippines

| Study Areas | Dengue Incidence | Allelic richness | Private allelic richness | Observed Heterozygosity | Inbreeding coefficient (Fis) | Median effective population size ($N_e$) |
| --- | --- | --- | --- | --- | --- | --- |
| CAL-N | 1.92 | 3.68 | 0.08 | 0.493 | 0.169 | 78.1 |
| CAL-S | 4.76 | 3.49 | 0.11 | 0.468 | 0.215 | 8.5 |
| VAL | 5.22 | 3.49 | 0.04 | 0.508 | 0.143 | 6.2 |
| QZC-1 | 0.25 | 3.24 | 0.10 | 0.470 | 0.111 | 116.1 |
| QZC-2 | 3.92 | 3.45 | 0.06 | 0.520 | 0.047 | 20.7 |
| QZC-3 | 3.26 | 3.44 | 0.03 | 0.525 | 0.071 | 57.7 |
| QZC-4 | 3.68 | 3.63 | 0.08 | 0.541 | 0.077 | 86.8 |
| QZC-5 | 4.45 | 3.37 | 0.05 | 0.500 | 0.117 | 58.2 |
| MNL-1 | 1.32 | 3.53 | 0.08 | 0.524 | 0.090 | 186.7 |
| MNL-2 | 3.70 | 3.55 | 0.03 | 0.468 | 0.132 | 386.8 |
| MRK | 3.64 | 3.85 | 0.07 | 0.511 | 0.112 | 4607.1 |
| SJ-MND | 4.08 | 3.29 | 0.03 | 0.496 | 0.191 | 38.1 |
| PSG | 2.81 | 3.56 | 0.08 | 0.524 | 0.099 | 31.7 |
| MKT | 2.02 | 3.59 | 0.06 | 0.558 | 0.036 | ∞ |
| PSY | 3.18 | 3.71 | 0.23 | 0.506 | 0.080 | 477.3 |
| TGG | 3.80 | 3.27 | 0.05 | 0.536 | 0.071 | 24.1 |
| PRNQ | 8.74 | 3.78 | 0.04 | 0.564 | 0.076 | 84.7 |
| LSP-1 | 8.37 | 3.65 | 0.12 | 0.536 | 0.104 | 38.4 |
| LSP-2 | 8.95 | 3.44 | 0.02 | 0.432 | 0.296 | ∞ |
| MTLP-1 | 8.44 | 3.79 | 0.13 | 0.438 | 0.240 | 79.4 |
| MTLP-2 | 7.51 | 3.51 | 0.03 | 0.449 | 0.240 | 94 |

### Genetic differentiation and structure

The overall *F*_ST_ was estimated to be 0.016 and pairwise *F*_ST_ values between study areas ranged from −0.002 – 0.054 (Table S6). With this, significant genetic differentiation was demonstrated in 87 (out of 201, 41.4%) pairwise values. Pairwise gene flow (Nm) estimates among study areas ranged from 3.404 – 290.448. 11 pairwise gene flow estimates were not calculated due to the estimated negative and zero *F*_ST_ values. The dendrogram based on the pairwise *F*_ST_ values revealed the spatial pattern and distribution of genetically similar study areas (Figure 2). Further analysis showed that the optimal number of cluster groups is 6 where 4 indicated groups of genetically similar study areas. It is shown that highly genetic-similar study areas are proximal to each other. This is exemplified by genetic group 2 where mosquito populations in the south area except for LSP-2 are genetically similar. There are also neighboring study areas that are genetically similar such as in genetic groups 3 (VAL, CAL-S), 1 (QZC-2 and QZC-4) and 4 (PSG-MRK). Furthermore, the pattern of the identified genetic groups extends in long distances as demonstrated in genetic groups 1 and 4.

**Figure 2.**
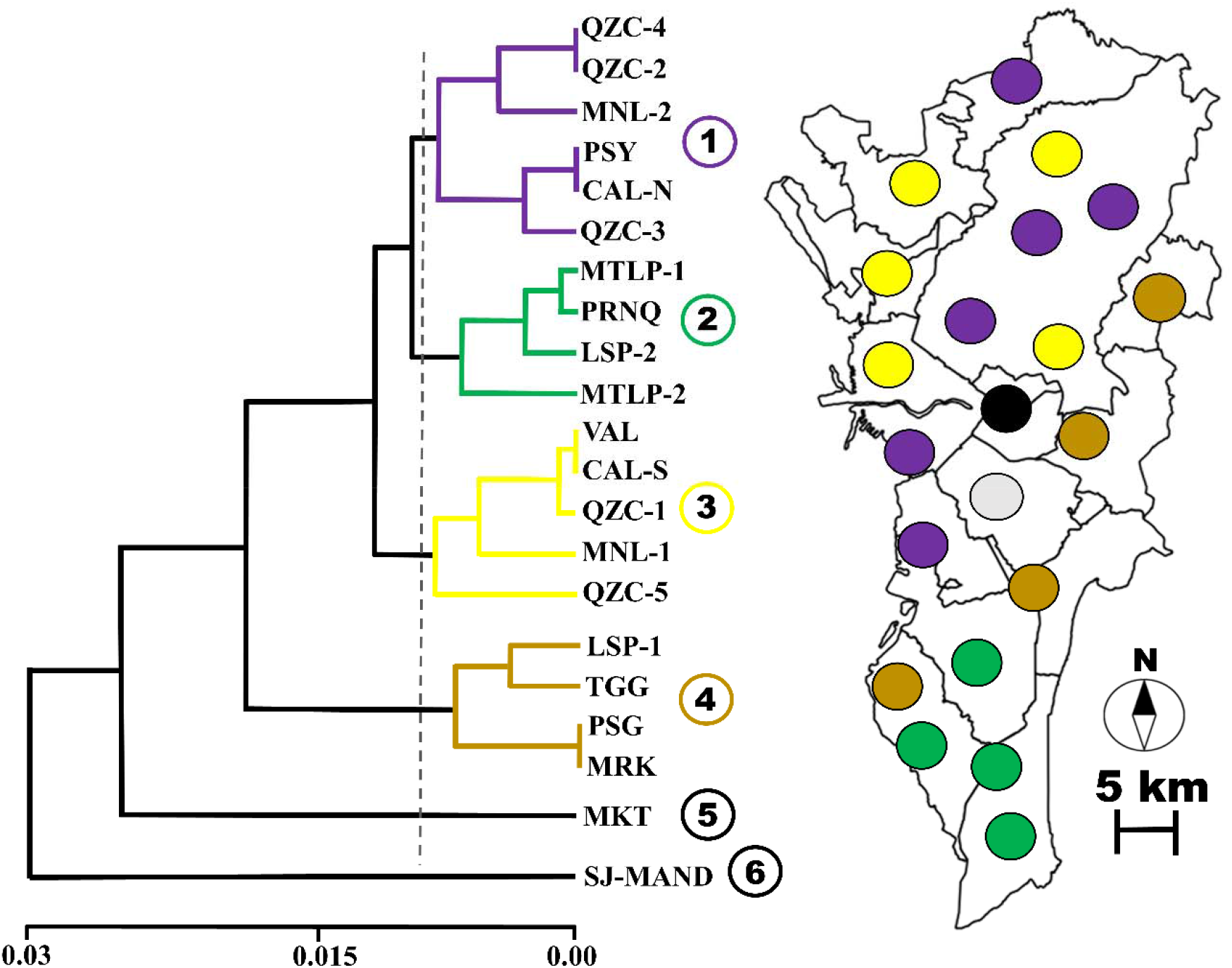
(a) Dendrogram showing the genetic relatedness of each study area based on its pairwise *F*_ST_ estimates. Colored lines indicate the genetic groups. (b) Map showing selected study areas in respect to their genetic group assignment. Colored circles indicate the genetic group.

Mantel test between the pairwise genetic (*F*_ST_) and geographic distances of all study areas showed very low and non-significant correlation (R^2^= 0.01, p=0.172), indicating no isolation by distance. High genetic similarity among adult mosquitoes is limited up to 1 km based on the spatial autocorrelation analysis (Figure S1). This suggests the limited dispersal capability of *Ae. aegypti*.

STRUCTURE analysis found that the most likely number of genetically differentiated groups is K = 5 (Figure S2). Figure 3 shows the distribution and proportion of inferred genetic cluster assignment of each mosquito individuals per study area. It is observed that either 4 or all inferred genetic clusters are present in each study area.

**Figure 3.**
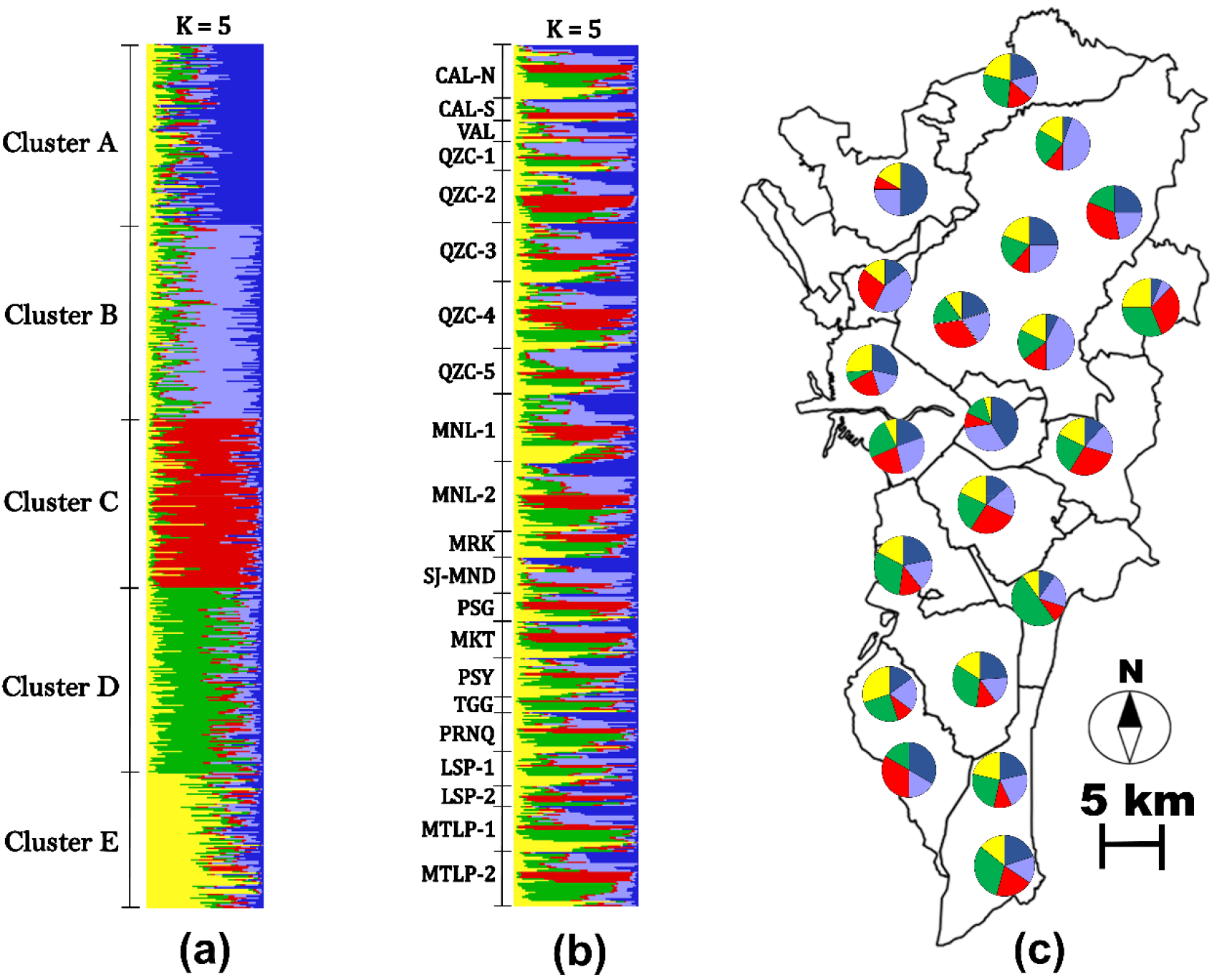
Bayesian analysis (K=5) of *Ae aegypti* populations in Metropolitan Manila. Bar plots represent the (a) genetic clusters and (b) study areas. Each individual is represented by a single horizontal line. Brackets are shown to separate genetic cluster or study areas. (c) Spatial map that shows the proportion of all genetic clusters to each study area. Colors represent the estimated individual proportion of cluster membership.

### Correlation between Population Genetic indices and Dengue incidence

Five genetic indices were used to correlate with dengue incidence. It revealed that allelic richness (*r = 0.30*) and Fis (*r = 0.52*) showed a positive correlation to dengue incidence. Among these indices, it was only Fis that showed statistical significance (Figure 4). On the other hand, private allelic richness (*r = −0.14*), observed heterozygosity (*r = −0.26*) and *Ne* (*r = −0.10*) demonstrated a negative correlation with no statistical significance.

**Figure 4.**
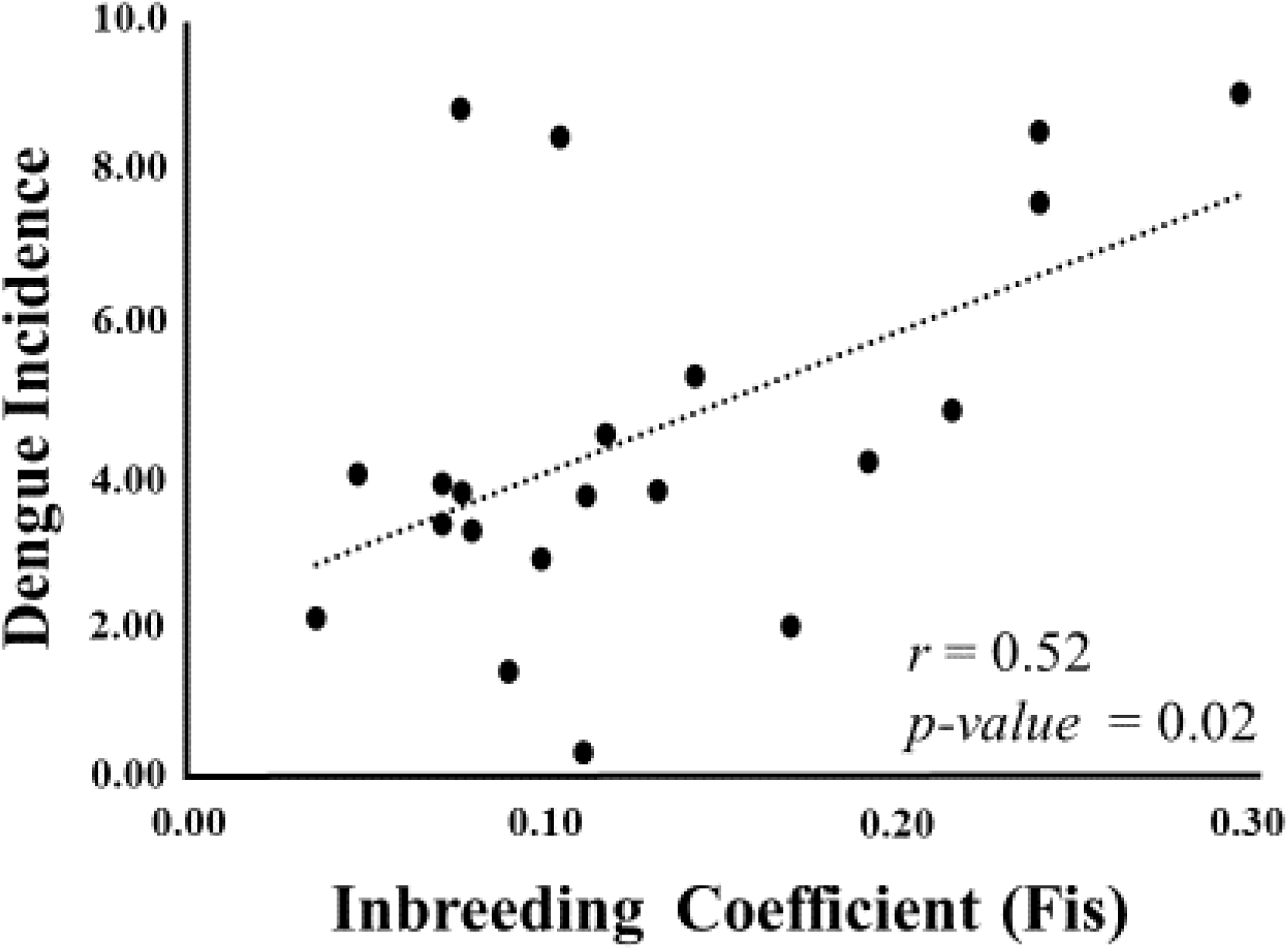
Correlation of Dengue Incidence and Inbreeding coefficient (Fis) among 21 study areas

## DISCUSSION

### Fine-Scale Genetic Structuring and Dispersal

Our study revealed low genetic differentiation among study areas (*F*_ST_ = 0.006 – 0.054) which is similar from previous population genetic studies of *Ae. aegypti* that consisted of a micro-geographic or fine-scale study area. In Sao Paulo, Brazil, for example, the level of genetic differentiation ranged from 0.002 to 0.094 with a maximum distance among collection sites of 30 km [14]. Cities in Southeast Asian countries also showed low levels of genetic differentiation from 0.026 - 0.032 with a spatial scale of 5 - 50 km [11]. The same was observed among villages in Thailand with geographical distances up to 27 km showing an average genetic differentiation of 0.037 [13]. Our findings along with previous studies suggest that mosquito populations within fine-scale areas may consist of similar allele frequencies, thus, exhibit continuous and active exchange or sharing of alleles among study areas. This is corroborated with the high gene flow estimates and the lack of a detected signal of isolation by distance observed in the study.

Our constructed dendrogram showed that genetically similar *Ae. aegypti* mosquito populations are in close proximity with each other. This clustering pattern is highly exemplified in the southern cities which comprises genetic group 2 (MTLP1, MTLP-2, PRNQ, LSP-2) and the eastern part of the region (MRK, PSG and TGG). It demonstrates that the dispersal of *Ae. aegypti* is limited which is also supported by our spatial autocorrelation analysis. We can only infer such limited dispersal could be attributed by landscapes in relation to the accessibility and location of each cities. For example, the southern cities which comprises genetic group 2 (MTLP1, MTLP-2, PRNQ, LSP-2) can only be accessed by one major highway while majority of the total land size of eastern cities (MRK, PSG and TGG) are completely separated by a major highway and a river (e.g. Marikina River). Therefore, access to these southern or eastern cities, may be difficult since a few roads, highways, or bridges connect it from the rest of the region. For this reason, it can potentially limit (but not isolate) the continuous migration and genetic exchange of mosquito populations from other study areas in the entire region. This information provides the potential migration or dispersal activity of *Ae. aegypti* that can be utilized in defining specific vector control zones along landscape corridors (e.g. roads) within certain group of cities.

What is also notable is that genetically-similar groups can extend in long distances such as observed in the northern to the central (genetic group 1) and the eastern to the southern parts (genetic group 4) of the region. If one overlays the major highways and road networks of Metropolitan Manila, it suggests the “passive” dispersal capability of *Ae. aegypti* by human-mediated transportation. It is believed that mosquito vectors occasionally travel in long distances by taking advantage human-aided transportation routes via land, sea or air [52,53,54] as *Ae. aegypti* eggs, larvae and adults have been found in commercial trucks and ships through tire importation [55,56]. In addition, transportation zones such as airports [57] and docks/ports [54,56] can be littered with larvae and pupae of the mosquito vector, thus acting as the source population. Rapid urbanization (e.g. commercialization) may also intensify this long distance migration or passive dispersal of *Ae. aegypti* by promoting mosquito population admixture over distant areas [15,58].

Due to the high genetic similarity and migration activity observed among study areas, we expected a low number of inferred genetic clusters (K=2 at the most). However, it revealed a considerable number of genetic clusters (K=5). These findings are consistent with previous population genetic studies with micro-geographic scales [11,14,16] and local studies in the Philippines [54,59]. Furthermore, these studies reported low genetic differentiation but with substantial number of inferred genetic clusters (K = 3-9). The substantial number of genetic clusters in fine-scale areas may be explained by two hypotheses. The first hypothesis could be the result of divergence from a single ancestry and, over time, produced multiple genetic clusters in this area. It is argued that a single or closely-situated houses may act as a clustering unit in forming genetically structured clusters [11,52,60]. This could be due to the limited flight performance of *Ae. aegypti* where it prefers to stay within a small area of about 10 – 500 meters [61] for a stable breeding site and availability of blood hosts [52]. As a result of rapid urbanization, these distinct genetic mosquito groups may have migrated to distant locations through “passive” dispersal, establishing colonies thereafter.

To some extent, our results support the first hypothesis where it revealed the limited dispersal capability of *Ae. aegypti* in short distances (up to 1 km) based from spatial autocorrelation analysis and, in turn, may have generated the 5 inferred genetically structured groups. Since Metropolitan Manila is considered highly urbanized with numerous transportation routes, it could have facilitated the passive dispersal of the mosquito vector in distant locations based on the interpretation of high gene flow activity among the study areas. However, limited dispersal cannot only be the single factor that can explain the occurrence of multiple clusters. Without high mutation, the mosquito population cannot diversify within a small spatial scale for a short evolutionary time. Hence, the second hypothesis infers that it could be due to the immigration of mosquito populations from neighboring regions or provinces of Metropolitan Manila, but a larger spatial scale genetic data is needed to test this hypothesis.

### Correlation of Genetic indices towards Dengue incidence

Among the genetic indices, we expected the estimated population size (*Ne*) to show a significant and positive correlation. However, it resulted in a non-significant and negative relationship with dengue incidence. In our study, collecting adult mosquito samples was the ideal choice to perform a better estimation of the effective population size (*Ne*). Previous population genetic studies conducted their sampling using either only larval samples or larval samples reared to adult stages but majority of mosquito larvae do not become adults in the natural setting [62]. Thus, collecting larvae to estimate the effective population size of the adult may be deemed inappropriate.

One possible reason of detecting no correlation between *Ne* and dengue incidence may be that the calculated *Ne* is either under- or over-estimated. Estimating the precise *Ne* of natural populations can be difficult, especially if there is a large time interval between sampling points or temporal disruptions such as migration or population replacement. It has been demonstrated that lower *Ne* estimates are generated by the presence of temporal disruptions while large intervals in sampling points generate higher *Ne* estimates [23]. Nevertheless, the calculated *Ne* estimates of the study is consistent with those calculated by Saarman et al. [23] from mosquito populations worldwide. But further and thorough investigation should be performed in the future to correctly estimate the *Ne* and its correlation with dengue incidence.

Notably, the correlation between dengue incidence and the inbreeding coefficient (Fis) revealed a positive and significant correlation (r =0.52, p =0.02). One plausible mechanism may be due to the transovarial or vertical transmission of the dengue virus to its succeeding mosquito offspring. This viral transmission process has been demonstrated in field-collected *Ae. aegypti* mosquitoes until its F2 generation from Quezon City, Metropolitan Manila [63]. Earlier studies also revealed that the transovarial route can sustain the viral infection up to the 15^th^ generation [64]. Therefore, we infer that the inbreeding within the mosquito population may result in producing a substantial number of next generation mosquitoes carrying the dengue virus. This suggests that dengue virus infection is not only maintained by the human-vector cycle but also within mosquito generations. Because of such mechanism, it can possibly lead to an increased disease transmission within local areas. It should be known there are limitations in our interpretation of this correlation. First, no viral detection was done to each individual *Ae. aegypti* mosquitoes to ascertain their viral transmission capability. Performing this endeavor could provide the necessary credence that can strongly support such correlation. Secondly, it is unclear whether the dengue virus infection in the human population originated from the exact study area. It has been suggested that dengue infections in the human population especially in urbanized areas can be obtained from other locations such as public spaces, schools or workplaces rather than their place of residence [65,66,67,68].

Nonetheless, this is the first attempt to directly link mosquito population genetic data to epidemiological data. Although several previous studies signified the application of population genetics towards vector control, they only reported the mosquito gene flow pattern and the association of mosquito genetic structure and its landscape [54,69,70]. Our findings may provide future implications in predicting or identifying high dengue risk areas where the genetic index (e.g. *F*is) can be utilized as a supplementary index with conventional mosquito-based indices. But further research is needed to ascertain on how these genetic indices can directly influence the local dengue epidemiology.

## SUPPORTING INFORMATION

**Figure S1.**
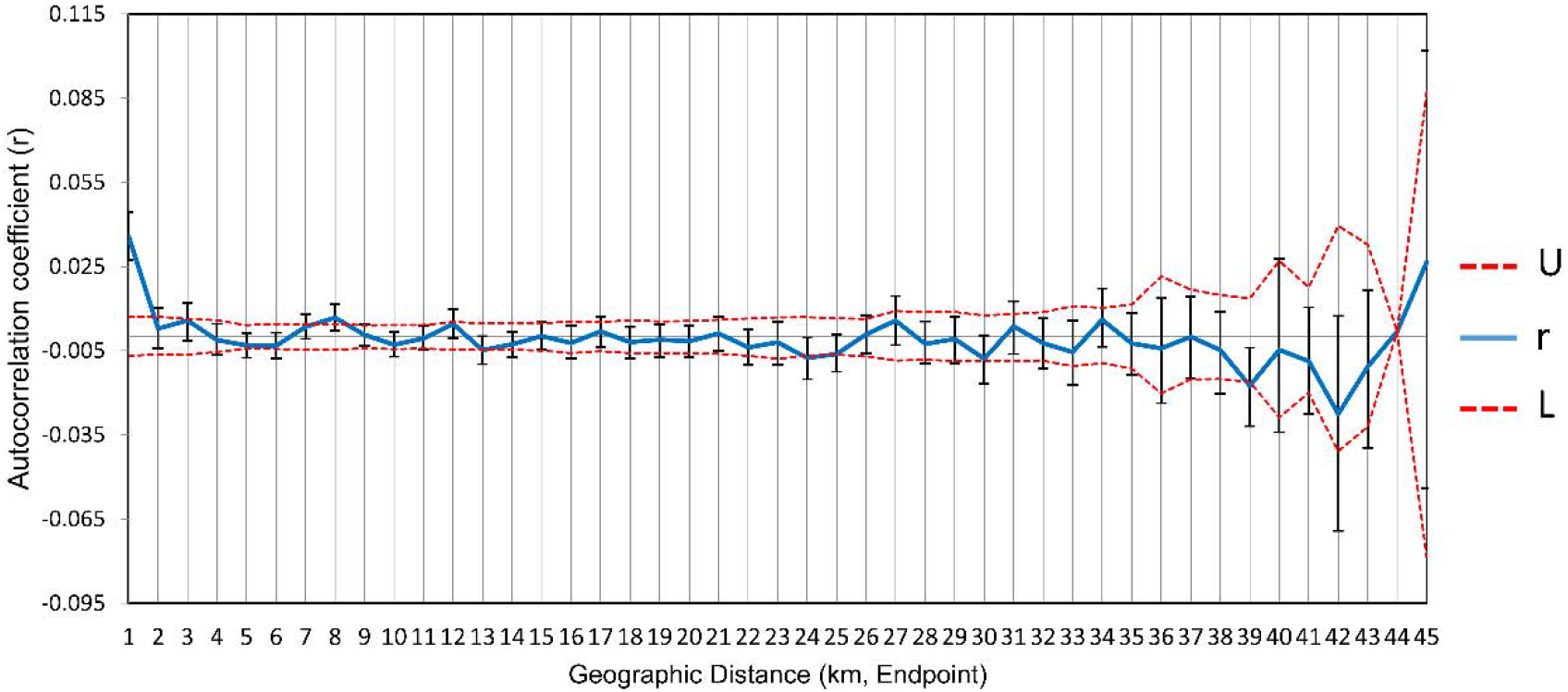
Correlogram of spatial autocorrelation showing the coefficient (r) up to 45 km with 1km intervals. U and L are upper and lower confidence interval limit respectively.

**Figure S2.**
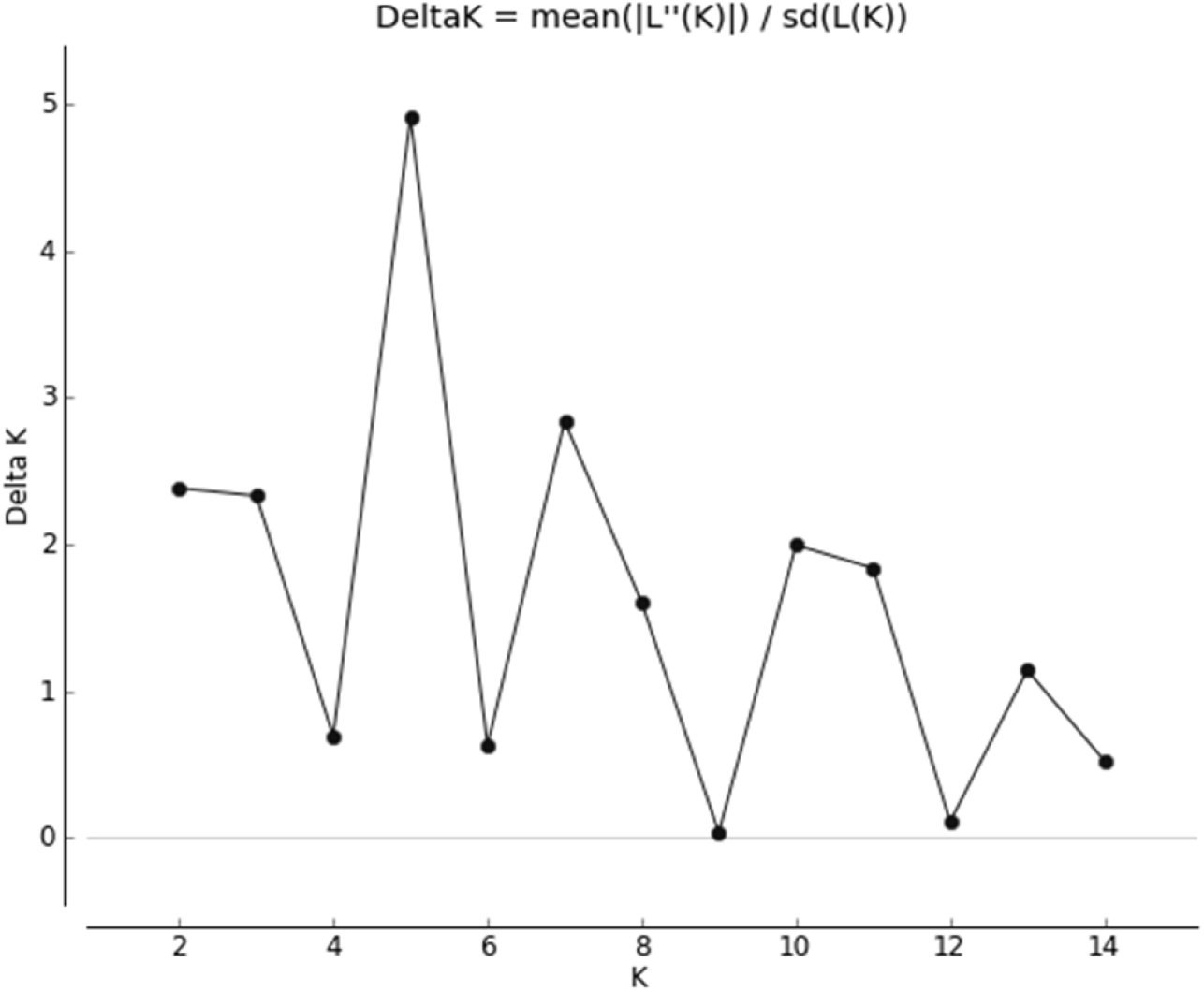
Graph of ΔK against K showing K = 5 as the probable number of genetic clusters

**Table S1.** *Ae aegypti* collection sites and population size in Metropolitan Manila, Philippines

**Table S2.** List and Characteristics of Microsatellites markers in *Ae. aegypti* used in this study

**Table S3.** Analysis of the Genetic Diversity of *Ae aegypti* using 11 microsatellite loci

**Table S4.** Null allele frequency estimates per locus per *Ae aegypti* populations

**Table S5.** Allele frequencies of the 11 microsatellite loci in *Ae aegypti* populations in Metropolitan Manila

**Table S6.** Genetic differentiation using *F*_ST_ (below diagonal) and Gene flow (*Nm*) (above diagonal) of *Ae aegypti* populations in Metropolitan Manila, Philippines

